# Inducing rapid seed germination of native cool season grasses with solid matrix priming and seed extrusion technology

**DOI:** 10.1101/413401

**Authors:** Matthew D. Madsen, Lauren Svejcar, Janae Radke, April Hulet

## Abstract

There is a need to develop effective techniques for establishing native vegetation in dryland ecosystems. We developed a novel treatment that primes seeds in a matrix of absorbent materials and bio-stimulants and then forms the mixture into pods for planting. In the development process, we determined optimal conditions for priming seeds and then compared seedling emergence from non-treated seeds, non-primed-seed pods, and primed-seed pods. Emergence trials were conducted on soils collected from a hillslope and ridgetop location on the Kaibab Plateau, Arizona, USA *Poa fendleriana* and *Pseudoroegneria spicata* were used as test species. Seeds were primed from −0.5 to −2.5 MPa for up to 12 d. Seeds primed under drier conditions (−1.5 to −2.5 MPa) tended to have quicker germination. Days to 50% emergence for primed-seed pods was between 66.2 to 82.4% faster (5.2 to 14.5 d fewer) than non-treated seeds. Seedling emergence from primed-seed pods for *P. fendleriana* was 3.8-fold higher than non-treated seeds on the ridgetop soil, but no difference was found on the other soil. Final density of *P. spicata* primed-seed pods were 2.9 to 3.8-fold higher than non-treated seeds. Overall, primed-seed pods show promise for enhancing germination and seedling emergence, which could aid in native plant establishment.

## Introduction

Nearly two-thirds of the globe’s ecosystems are considered degraded [1]. As an example, in North America, the sagebrush biome is declining at an alarming rate as large-scale catastrophic wildfires and other disturbances remove native vegetation. These losses allow for the invasion of exotic annual grasses such as *Bromus tectorum* (downy brome or cheatgrass) [2, 3]. Exotic annual grass invasion further promotes the frequency, extent, and severity of wildfires, which creates more disturbed areas for exotic annual grasses to colonize [4]. Effective methods for seeding native vegetation back into degraded sagebrush systems are needed to prevent or reduce weed invasion and arrest the invasive plant-fire cycle. Successful seeding of native vegetation can also help to preserve soil and water resources, sustain wildlife habitat, increase forage production and enhance landscape aesthetics [5]. Unfortunately, success rates for reestablishing native plants from seeds in disturbed sagebrush systems and other dryland regions are unacceptably low [3, 6], and seeding success is predicted to further decline with climate changes, such as increasing aridity and more erratic precipitation [7].

Seed enhancement technologies, defined as treatments applied after seed harvest and before sowing that improve seed delivery to a system, seed germination or seedling growth [8], are being developed to improve seeding success in dryland systems [9-13]. Characteristics or mechanisms that allow non-native annual grasses to invade following disturbance may be key to guide and develop new seed enhancement technologies. A major trait associated with the spread and dominance of exotic annual grasses is their ability to rapidly germinate and emerge from the soil in high numbers several days earlier than native perennials [14, 15]. Early germination gives invasive annual weeds temporal priority for short term, limited moisture resources, allowing them to out-compete native perennial species attempting to establish from seed [15].

Priming is a seed technology that allows seed germination processes to begin by partially hydrating seeds at water potentials that allow imbibition [16]. Water uptake by seeds occurs in Phase 1 of germination and plateaus in Phase 2 until water uptake is again initiated by Phase 3 post-germination growth (i.e. when a part of the embryo, typically the radicle, grows to penetrate the seed exterior) [17]. Seed priming has been suggested as a possible treatment to provide native cool-season grasses with similar germination characteristics as exotic annual grasses [15, 18]. “Osmopriming” is a common priming approach that places seeds in an aerated osmotic solution to induce water stress [19]. Priming osmoticum commonly used include polyethylene glycol (PEG), inorganic salts or mannitol. The osmotic potential of the solution is adjusted to allow the seed to complete early phases of germination (e.g. Phase 1 and 2, prior to the priming treatment being arrested). Osmopriming has several technical and logistical difficulties associated with the practice [20]. Specialized equipment is required so osmotic solutions are continuously aerated. Also, viscosity and oxygen diffusivity problems can occur when osmotic solutions are too concentrated. Differences between priming osmoticum are also found. For example, inorganic salts can penetrate through the seed coat, adversely affecting seed germination. PEG is a large molecule that cannot penetrate into seed tissues, but it is relatively expensive, with large volumes of chemical solution required in relation to the quantity of seeds used. Further cost is incurred in the disposal of the product [20].

Solid matrix priming (SMP) mixes seeds with a solid carrier that is moistened with water to achieve desired water potentials for priming. There is evidence to suggest that this type of priming is as effective and in some cases, more effective than osmotic priming [20, 21]. The major limitation with SMP is that after seeds are primed, the solid matrix material needs to be mechanically separated without harming the seeds.

Challenges associated with SMP may be alleviated if seeds could be efficiently planted with the SMP medium. Furthermore, seeding efforts may be improved if the SMP material enhanced seed germination and seedling growth. Madsen and Svejcar [22] showed that seeds could be placed together in extruded pods using machinery for making pastas and pastries. Pods are formed by creating a “dough” containing seeds, various clay filler materials, absorbents, bio-stimulants, plant protectants, water, and other desired ingredients, and then running the mixture through an extruder that forms and cuts the extruded material into desired shapes. Seed pods are designed for broadcast seeding by providing seed coverage and enhanced conditions for seed germination and growth [22].

The purpose of this study was to develop methods for priming seeds using SMP techniques in the material used to form extruded-seed pods. Specific objectives were to: 1) assess the suitability of the seed-extrusion material as a priming medium and develop a moisture release curve, 2) determine optimal water potentials and priming durations for priming seeds in the dough, and 3) compare seedling emergence between non-treated seeds, non-primed-seed pods, and primed-seed pods on two different soil types.

## Materials and methods

### Experiment 1: Estimation of optimal-priming conditions

Our research was conducted on two perennial bunchgrasses ‘Ruin canyon’ muttongrass (*Poa fendleriana* (Steud.) Vasey) and ‘Anatone’ bluebunch wheatgrass (*Pseudoroegneria spicata* (Pursh) Löve). Both species are commonly used for rangeland restoration projects in the western United States. Previous work suggests *P. spicata* has a faster germination velocity than species of Poa native to the Great Basin [23].

The effect of water potential and seed priming duration on seed germination was assessed at water potentials of −0.5, −1.0, −1.5, −2.0, and −2.5 MPa for either 4, 5, 6, 7, 8, 9, 10, 11, or 12 days (d). The combination of differing water potentials and seed priming durations are designated as “priming periods”. Seeds were primed in a solid matrix containing calcium bentonite clay, diatomaceous earth, compost, worm castings, non-ionic alkyl terminated block co-polymer surfactant, plant growth regulator, fungicide, liquid fertilizer, and tap water (Table 1). Worm castings and compost were air-dried and screened through a 1 mm sieve. The amount of seed added to the recipe was equal to the amount required to produce approximately eight pure live seeds (PLS) in an extruded seed pod that was 20 mm long × 20 mm wide × 6 mm thick (see the following experiment for methods to produce extruded seed pods; Table 1). Twenty pods were checked for PLS after every preparation to guarantee 8 PLS per pod.

**Table 1.** Recipe for producing extruded seed pods under optimal priming conditions. The recipe was used on 8.11 and 32.48 g of ‘Ruin canyon’ *Poa fendleriana* and ‘Anatone’ *Pseudoroegneria spicata* seed, respectively. This amount of seed was needed to produce approximately 8 pure live seed pod^-1^.

| Ingredients |  |  |  |
| --- | --- | --- | --- |
| Trade name | Product | Supplier | Batch wt.<br>(g) |
| Pelbon | Ca Bentonite | American Colloid Co. (Arlington Heights, IL) | 557.6 |
| BioFine | Compost | Deschutes Recycling (Bend, OR) | 520.0 |
| DiaSource | DE<sup>†</sup> | DiaSource, Inc (Boise, ID) | 203.8 |
| Ignite | Fertilizer | Land View Inc. (Rupert, ID) | 4.6 |
| Captan | Fungicide | Arysta LifeScience (Cary, N.C.) | 2.5 |
| Ascend | PGR<sup>‡</sup> | Winfield Solutions (St Paul, MN) | 3.6 |
| Stockobsorb 660 granuals | SAP<sup>§</sup> | Evonik Stockhausen (Greensboro, NC) | 25.0 |
| Stockobsorb 660 powder | SAP | Evonik Stockhausen (Greensboro, NC) | 60.0 |
| Startch 1500 | Starch | Colorcon (West Point, PA) | 40.0 |
| ASET-4001 | Surfactant | Aquatrols Corporation (Paulsboro, NJ) | 1.6 |
| Worm Gold | Worm Castings | California Vermiculture (Cardiff, CA) | 200.0 |

| Priming water potential<br>(MPa) | Water content<br>(%) | Water required for priming<br>(g) | Additional water to produce seed pods<br>(g) |
| --- | --- | --- | --- |
| -0.5 | 50.8 | 781.4 | 1,349.0 |
| -1 | 38.5 | 589.6 | 1,540.7 |
| -1.5 | 30.9 | 472.4 | 1,656.5 |
| -2 | 26.9 | 408.8 | 1,721.5 |

<sup>†</sup> Diatomaceous earth
<sup>‡</sup> Plant Growth Regulators
<sup>§</sup> Superabsorbent polymers

To determine how much liquid was needed to achieve a desired water potential for priming, we developed a moisture release curve for the solid matrix priming medium by adjusting the moisture of the medium with water at approximately 2% intervals (expressed as a percent dry weight) from 12 – 100%. At each moisture interval, water potential of the medium was measured with a WP-4 Dewpoint Potentiameter (Decagon Devices, Inc., Pullman, Washington, USA). CurveExpert 1.4 (Hyams Development, USA) was used to fit a nonlinear regression equation through the data based on maximum R^2^ and F-values with minimum residuals (Fig. 1). This nonlinear regression equation was then used to estimate the amount of water required to achieve the desired water potential in the priming medium (Fig. 1; Table 1).

**Fig 1.**
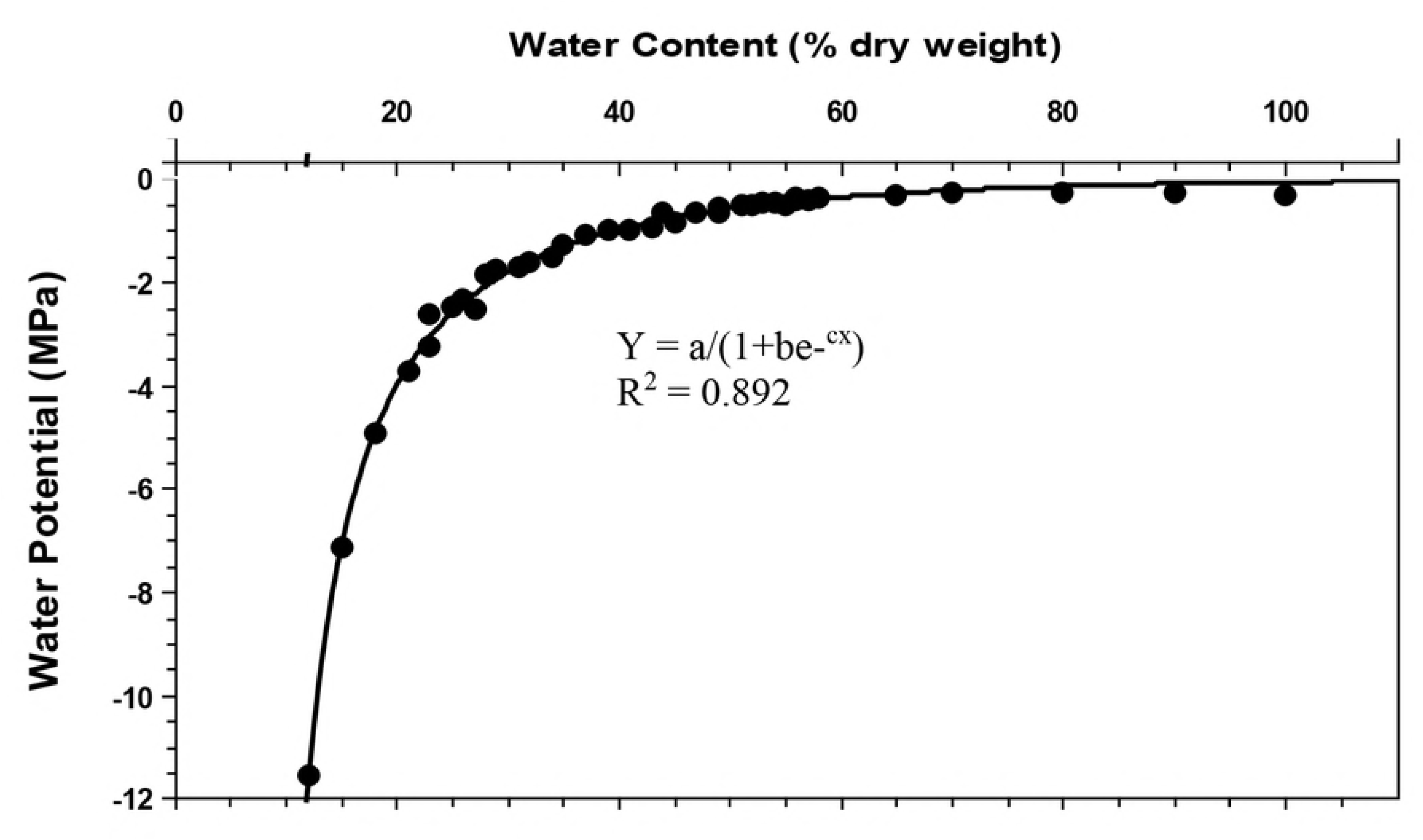
Moisture release curve of material used for producing extruded seed pods and pellets. Water content is expressed as percentage based on the dry weight of the solid material. CurveExpert 1.4 was used to fit curve; best fit curve used a logistic model where a = 3.1441, b = −7.5929E-001, and c = −4.2990E-002.

Surfactant, plant growth regulator, fungicide, liquid fertilizer, and tap water were combined into a homogeneous mixture when preparing the priming medium for each unique water potential/priming duration combination and for each species. This liquid solution was then applied to a mixture of seeds, calcium-bentonite clay, diatomaceous earth, compost, and worm castings using a 1 L KitchenAid mixer (Joseph, MI, USA) with a wire whip. After mixing for 3 minutes (min) the material was transferred in equal amounts into three separate 5.7 L box storage containers (33 cm length × 20 cm width × 13 cm depth). Priming boxes were incubated in a SG50SS environmental growth chamber (Hoffman Manufacturing Inc., Jefferson, OR) at a constant temperature of 15 ± 0.5 °C and a daily light period of 12 hd^-1^ with fluorescent lights.

Starting on day four of the priming process, we weighed every sample each day and replaced the water lost by evaporation. Material in the priming boxes was also lightly mixed by hand each day and seeds were examined for coleorhizae protrusion from a 30 g subsample. This process was done to determine the duration that seeds could be incubated in the priming medium without completing germination (i.e. post-germination radical emergence). After each of the specified priming periods, seeds were extracted from the priming medium by sieving. Seeds were then air dried on a laboratory benchtop. Water potential/priming duration combinations having >10% of the seeds with coleorhiza protruding from the seeds were discarded from the study.

To assess changes in percent germination, four sets of 25 seeds were evaluated for each unique priming period treatment in 8 cm diameter Petri dishes that contained a single layer of blue blotter paper (Anchor Paper Co., St. Paul, Minn.). The study was arranged in a randomized complete-block design. Petri dishes were placed in a SG50SS environmental growth chamber (Hoffman Manufacturing Inc., Jefferson, OR) and incubated at 10 ± 0.5 °C. Seed germination was counted twice daily (in the morning and afternoon) for the first 16 days and then once daily for 9 days (25 days total); seeds were considered germinated when a radicle extended at least 2mm out of the seed. Petri dishes were rotated on different shelves in the growth chamber throughout the experiment. The growth chamber was set for a 12 hd^-1^light period with fluorescent lights.

From daily germination counts, we calculated the following germination indices: 1) mean germination time (MGT), time to reach 10, 50, and 90% germination (T_10g_, T_50g_, T_90g_), germination synchrony, and final germination percentage (FGP).

MGT was calculated according to the following equation:

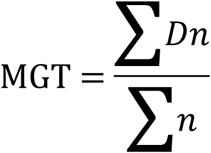

where:

n=The number of seeds that germinated on day D

D=The number of days counted from the beginning of germination

Time to reach T_10g_, T_50g_, and T_90g_ was calculated as follows:

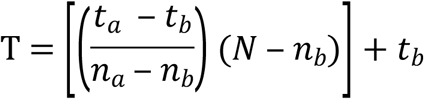

where:

T=time (days) to subpopulation germination

t_a_=incubation day when subpopulation germination was reached

t_b_=incubation day before subpopulation germination was reached

n_a_=number of germinated seeds on day that subpopulation germination was reached

n_b_=number of germinated seeds on day before subpopulation germination was reached

N=number of germinated seeds equal to 10, 50 or 90% of the total population

Germination synchrony was estimated by subtracting T_90g_ from T_10g_ (i.e., T_90g_-T_10g_). Final germination percentage (FGP) was calculated as the ratio of the number of seeds germinated to the total number of seeds evaluated and was expressed as a percentage. The priming treatment that produced the quickest germination timing (based off of T_50g_ values) without impacting FGP was used in Experiment 2.

### Experiment 2: Evaluation of primed extruded seed pods

Seedling emergence was compared between non-treated seeds (control), non-treated-seed pods, and primed-seed pods with *P. fendleriana* and *P. spicata* seeds. Separate evaluations were conducted for each species on two different soil types collected from the West Side region of the Kaibab Plateau, Arizona USA. Soils were obtained within 4 km of each other; one collection site was on a ridgetop (36° 37’ 28.41” N, 112° 31’ 25.15’ W), and the other from a hillslope (36° 38’ 51.69’ N, 112° 29’ 19.73’ W). Mean annual precipitation and temperature in this area was 360-440 mm and 7-9 °C, respectively [24]. At both collection sites, soils were derived from Kaibab limestone, which is comprised of a diverse assemblage of sedimentary rock types. At the ridgetop, the soil was less developed relative to the hillslope site and had a high quantity of lime throughout the soil profile [24]. Soil texture at the ridgetop site was a gravelly sandy-loam and classified as a Lithic Ustochrepts, calcareous loamy-skeletal, mixed mesic [24] with a soil pH of 7.5. At the hillslope site, soil texture was a very gravely fine sandy-loam and classified as a Lithic Ustochrepts, calcareous, loamy-skeletal, mixed, mesic [24] with a soil pH of 7.4. The two soil types were selected because of observed differences in restoration success by management groups and demonstrated differences in water holding capacity.

Materials for producing seed pods were the same as used in Experiment 1, with the exception that the recipe also included synthetic superabsorbent polymer fine granules and powder (Table 1). It is important that the medium used be capable of providing a constant water potential throughout the priming duration [20]. Superabsorbent polymers were not included during priming in Experiment 1 because in a preliminary study it was observed that a constant water potential could not be maintained. Observations revealed that the superabsorbent polymers would hydrate when water was added but overtime the absorbed water would be released back into the surrounding medium, thus increasing the water potential.

To produce seed pods, ingredients were mixed into a dough using a 1 L KitchenAid mixer (Joseph, MI) for a period of 3 min. The seed dough was then extruded and cut by hand into 20 mm long × 20 mm wide × 6 mm thick pods with eight PLS per pod [22]. After cutting, pods were dried on a forced air dyer at 43°C. To produce primed extruded seed pods, *P. fendleriana* seeds were primed for 10 d at −1.5 MPa; *P. spicata* seeds were primed for 6 d at −2.0 MPa, as described in Experiment 1. Immediately after seeds were primed, superabsorbent polymers were mixed into the medium. Additional water was then added to the seeds and priming material (Table 1) and mixed into a dough, formed into pods, and dried following the steps used to create non-primed-seed pods.

Each soil type was compacted into two 16 L wooden boxes (50 cm length × 40 cm width × 8 cm depth). One of the two boxes for each soil type was seeded with *P. fendleriana;* the other box was seeded with *P. spicata.* Within a box, we compared the following seed treatments: 1) non-treated seeds, 2) non-primed-seed pods, and 3) primed-seed pods in a randomized-complete block design with six replicates (rows) per treatment. Seeds of each treatment were sown on the soil surface in 20 cm rows. Each row contained three pods, which equaled 24 PLS per row.

Soil boxes were placed in an environmental grow-room set at a constant temperature of 21°C, and a 12 hd^-1^ light period with 632 W m^-2^ fluorescent lighting. Prior to seeding, the soil was watered with a fine mist sprayer to 50% of field capacity, as determined using the “container capacity” method [25]. Following planting, 1 cm of water was applied (2 L of water per box). Over the remainder of the study, boxes were watered weekly with 1 cm of water. Plant density was measured for each row every 1-2 days for 31 days. From emergence counts, we determined the time to reach 50% emergence (T_50e_), mean emergence time (MET), and final emergence (FE) as described in Experiment 1.

Data was analyzed in SAS Version 9.4 (SAS Institute Inc., Cary, NC, USA). We used mixed model analysis to analyze T_50e_, MET, and FE. In the model, seed treatments were considered a fixed factor and blocks a random factor. Prior to analysis, data were tested for normality using the Shapiro Wilk test. Where appropriate data were log or square root transformed prior to analysis to achieve normal distribution. When significant effects were found, mean values were separated using the LSMEANS procedure, with P-values adjusted using a Tukey test (P<0.05). Original (i.e., non-transformed) data are presented in the text and figures. We graphically displayed seedling density over the period of the study for each treatment by species. Plant density was calculated by dividing the number of plants in the row by the row length and width. The row width was assumed to be 30.5 cm, which represents a typical seeding width produced by a rangeland seed drill.

## Results

### Experiment 1: Estimation of optimal-priming conditions

The duration seeds could be primed without causing germination in the priming medium varied with water potential and between species. *Poa fendleriana* had a wide range in which seeds could be primed wherein it did not show coleorhizae protrusion at any water potential (−0.5, −1.0,−1.5, −2.0, or −2.5 MPa) for any time period (4, 5, 6, 7, 8, 9, 10, 11, or 12 d). However, *P. spicata* had a relatively narrow range of water potentials (−1.5, −2.0, and −2.5 MPa) and durations (4-7 d) in that seeds could be primed without showing coleorhizae protrusion.

Primed seeds of *P. fendleriana* germinated substantially faster than non-primed seeds at all water potentials and priming durations tested (Table 2). A general trend was observed where germination velocity would increase with priming duration. However, a threshold was identified for each water potential-priming duration treatment past which point either improvement halted (e.g. −2.5 MPa for 10, 11 and 12 d had similar germination velocities, Table 2) or seeds would germinate during priming (e.g. −1.5 MPa for 8 d had over 10% radicle emergence, Table 2). Priming seeds in the drier water potentials (−1.5, −2.0, and −2.5 MPa) for longer periods of time tended to produce quicker germination. Seeds primed for 10 d at −1.5 MPa had the quickest germination response. At this water potential and duration, priming decreased T_50g_ by 66.6% (10 d) and MGT by 61.2% (9 d). The synchrony of germination also improved with decreasing germination times (Table 2). Germination of *P. fendleriana* seeds primed for 10 d at −1.5 MPa were most synchronous, with T_50g_-T_10g_ values 64.9% (10.5 d) less than non-primed seeds. In general, most water potentials and priming durations had a slight improvement in the number of seeds that germinated in comparison to non-treated seeds (Table 2). Drier water potentials (−1.5,−2.0, and -2.5 MPa) tended to have a higher FGP after being primed for 10 d or less. Seeds primed for 10 d at −1.5, −2.0, and −2.5 MPa were similar to each other and had a FGP that was 20% higher than non-primed seeds. Based on the results of this study, priming at −1.5 MPa for 10 d appears to be optimal for *P. fendleriana*; however, priming at −2.0 and −2.5 MPa for 10-11 d may also be effective.

**Table 2.** Influence of matrix potential (Ψ) and priming duration on *Poa fendleriana* germination.

| Indices | $\Psi$ | Priming duration<br>(days) | | | | | | | | | |
| --- | --- | --- | --- | --- | --- | --- | --- | --- | --- | --- | --- |
|  |  | 0 | 4 | 5 | 6 | 7 | 8 | 9 | 10 | 11 | 12 |
| $T_{50g}^{\dagger}$ (days) | 0 | 14.3 (0.2) | | | | | | | | | |
|  | -0.5 |  | 10.8 (0.1) | 9.5 (0.5) | 8.2 (0.2) | 7.7 (0.2) |  |  |  |  |  |
|  | -1 |  | 10.7 (0.0) | 8.4 (0.0) | 7.3 (0.2) | 7.4 (0.3) | 6.7 (0.6) |  |  |  |  |
|  | -1.5 |  | 10.2 (0.2) | 9.2 (0.3) | 8.1 (0.3) | 7 (0.1) | 6.5 (0.3) | 5.8 (0.2) | 4.8 (0.1) | 5.4 (0.1) |  |
|  | -2 |  | 11.3 (0.3) | 10.3 (0.1) | 8.9 (0.3) | 7.6 (0.1) | 7.3 (0.2) | 6.1 (0.3) | 5.5 (0.4) | 5.9 (0.3) | 5.6 (0.4) |
|  | -2.5 |  | 10.6 (0.1) | 10.2 (0.2) | 9.7 (0) | 9.2 (0.6) | 8.3 (0.4) | 7.3 (0.2) | 6.1 (0.4) | 6.1 (0.2) | 6.2 (0.2) |
| $MGT^{\ddagger}$ (days) | 0 | 14.3 (0.1) | | | | | | | | | |
|  | -0.5 |  | 11.5 (0.4) | 10.3 (0.4) | 9.8 (0.6) | 8.3 (0.1) |  |  |  |  |  |
|  | -1 |  | 11.4 (0.1) | 9.8 (0.1) | 8.2 (0.1) | 8.5 (0.5) | 7.6 (0.3) |  |  |  |  |
|  | -1.5 |  | 11.0 (0.2) | 9.9 (0.1) | 9.3 (0.2) | 7.9 (0.3) | 7.4 (0.4) | 7.1 (0.3) | 5.6 (0.2) | 6.4 (0.3) |  |
|  | -2 |  | 12.1 (0.3) | 11.2 (0.2) | 10.3 (0.4) | 8.7 (0.1) | 8.1 (0.2) | 7.4 (0.4) | 6.9 (0.3) | 6.5 (0.3) | 6.5 (0.2) |
|  | -2.5 |  | 11.2 (0.2) | 11.1 (0.3) | 10.5 (0.1) | 10 (0.3) | 9.2 (0.1) | 8.3 (0.2) | 7.4 (0.2) | 7.0 (0.2) | 7.0 (0.2) |
| $T_{90g} - T_{10g}^{\S}$ (days) | 0 | 16.2 (1.1) | | | | | | | | | |
|  | -0.5 |  | 13.3 (0.3) | 13.3 (1.2) | 11.6 (0.7) | 10.7 (3) |  |  |  |  |  |
|  | -1 |  | 12.6 (0.3) | 13.2 (1.1) | 9.3 (0.3) | 10.3 (1.3) | 10 (1.8) |  |  |  |  |
|  | -1.5 |  | 12.6 (1.4) | 10.7 (0.2) | 11 (1) | 9.1 (0.7) | 9.4 (0.9) | 7.4 (0.5) | 5.7 (0.2) | 8 (0.9) |  |
|  | -2 |  | 12.7 (0.3) | 11.9 (0.3) | 10.8 (0.7) | 9.3 (1) | 8.7 (0.5) | 8.2 (0.4) | 7.5 (0.7) | 8 (0.7) | 6.8 (0.8) |
|  | -2.5 |  | 12 (1) | 12.2 (0.8) | 10.9 (0.3) | 11.3 (1.4) | 9.4 (0.4) | 8.8 (0.4) | 8.4 (1.2) | 7.5 (0.4) | 8.1 (0.6) |
| FGP<sup>¶</sup> (%) | 0 | 69 (3) |  |  |  |  |  |  |  |  |  |
|  | -0.5 |  | 76 (4) | 69 (4) | 83 (3) | 68 (5) |  |  |  |  |  |
|  | -1 |  | 80 (2) | 78 (5) | 78 (5) | 75 (8) | 78 (6) |  |  |  |  |
|  | -1.5 |  | 84 (7) | 85 (3) | 82 (3) | 79 (3) | 78 (3) | 90 (3) | 83 (3) | 75 (1) |  |
|  | -2 |  | 85 (3) | 84 (4) | 92 (4) | 88 (3) | 83 (3) | 83 (2) | 87 (2) | 73 (2) | 76 (8) |
|  | -2.5 |  | 83 (4) | 82 (3) | 84 (3) | 85 (4) | 82 (8) | 88 (4) | 85 (4) | 86 (4) | 77 (6) |

<sup>†</sup> $T_{50g}$ = time to 50 percent germination
<sup>‡</sup>MGT = mean germination time
<sup>§</sup> $T_{90g} - T_{10g}$ = difference between time to 90% and 10% germination
<sup>¶</sup>FGP = final germination percentage
Numbers in parentheses represent s.e.m.

Priming of *P. spicata* seeds also decreased germination time at all tested water potentials and priming durations (Table 3). Trends in germination time and priming duration were less apparent due to the inherently faster germination time of *P. spicata* in comparison to *P. fendleriana*. Most priming durations and water potentials were similar to each other. On average, T_50g_ decreased with priming duration at −2.0 and −2.5 MPa. Seeds that were primed for 6 d at −2.0 MPa and 7 d at −2.5 MPa had T_50g_ values that were approximately 50% (5 days) less than non-primed seeds. Mean germination time response was similar to T_50g_. Priming typically did not influence T_50g_-T_10g_ or FGP indices for *P. spicata*.

**Table 3.**
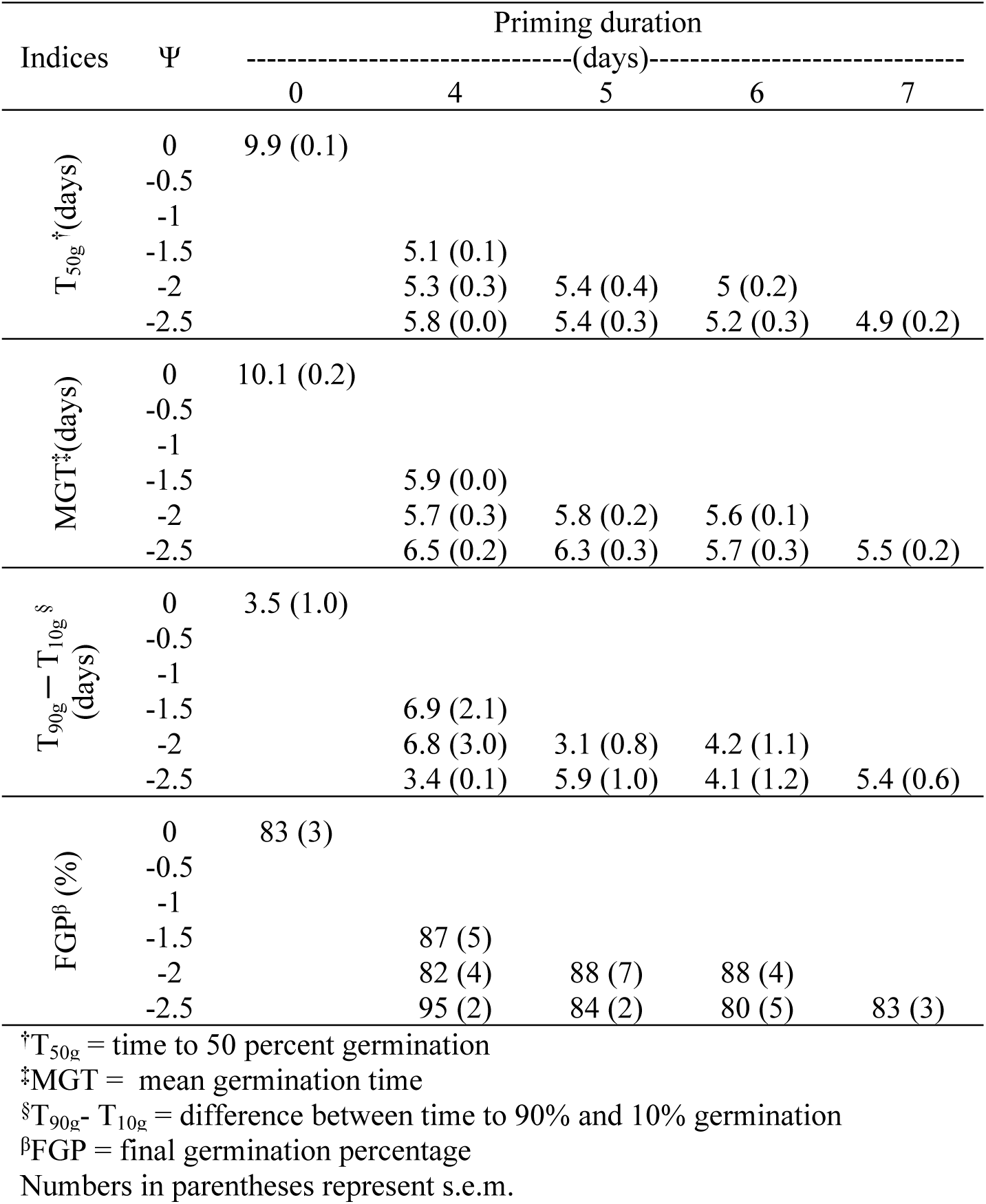
Influence of matrix potential (Ψ) and priming duration on *Pseudoroegneria spicata* germination.

| Indices | $\Psi$ | Priming duration | | | | |
| --- | --- | --- | --- | --- | --- | --- |
|  |  | (days) |  |  |  |  |
|  |  | 0 | 4 | 5 | 6 | 7 |
| $T_{50g}^{\dagger}$ (days) | 0 | 9.9 (0.1) | | | | |
|  | -0.5 |  |  |  |  |  |
|  | -1 |  |  |  |  |  |
|  | -1.5 |  | 5.1 (0.1) |  |  |  |
|  | -2 |  | 5.3 (0.3) | 5.4 (0.4) | 5 (0.2) |  |
|  | -2.5 |  | 5.8 (0.0) | 5.4 (0.3) | 5.2 (0.3) | 4.9 (0.2) |
| MGT$^{\ddagger}$ (days) | 0 | 10.1 (0.2) | | | | |
|  | -0.5 |  |  |  |  |  |
|  | -1 |  |  |  |  |  |
|  | -1.5 |  | 5.9 (0.0) |  |  |  |
|  | -2 |  | 5.7 (0.3) | 5.8 (0.2) | 5.6 (0.1) |  |
|  | -2.5 |  | 6.5 (0.2) | 6.3 (0.3) | 5.7 (0.3) | 5.5 (0.2) |
| $T_{90g}-T_{10g}^{\S}$ (days) | 0 | 3.5 (1.0) | | | | |
|  | -0.5 |  |  |  |  |  |
|  | -1 |  |  |  |  |  |
|  | -1.5 |  | 6.9 (2.1) |  |  |  |
|  | -2 |  | 6.8 (3.0) | 3.1 (0.8) | 4.2 (1.1) |  |
|  | -2.5 |  | 3.4 (0.1) | 5.9 (1.0) | 4.1 (1.2) | 5.4 (0.6) |
| FGP$^{\P}$ (%) | 0 | 83 (3) | | | | |
|  | -0.5 |  |  |  |  |  |
|  | -1 |  |  |  |  |  |
|  | -1.5 |  | 87 (5) |  |  |  |
|  | -2 |  | 82 (4) | 88 (7) | 88 (4) |  |
|  | -2.5 |  | 95 (2) | 84 (2) | 80 (5) | 83 (3) |

$^{\dagger}T_{50g}$ = time to 50 percent germination
$^{\ddagger}$ MGT = mean germination time
$^{\S}T_{90g}-T_{10g}$ = difference between time to 90% and 10% germination
$^{\P}$ FGP = final germination percentage
Numbers in parentheses represent s.e.m.

### Experiment 2: Evaluation of primed extruded seed pods

Seed treatments had a strong influence on T_50e_ for *P. fendleriana* and *P. spicata* in both soil types (*P. fendleriana* ridgetop *F* = 10.85, *P* < 0.001, hillslope *F* = 12.24, *P* < 0.001; *P. spicata* ridgetop *F* = 12.60, *P* < 0.001, hillslope *F* = 7.37, *P* = 0.01). Mean T_50e_ values from primed-seed pods were between 66.2-82.4% less than the control, depending on the species and soil type. Seedlings were recorded emerging from the primed-seed pods as early as 2 d after planting (Fig. 2). The majority of seedlings typically emerged from primed-seed pods within the first 10 d of the study. Seedling densities from non-primed-seed pods and the control took over 23 d or more to plateau (Fig. 2). Mean emergence times generally followed a similar pattern as T_50e_ (Table 4).

**Table 4.** Mean seedling emergence from *Poa fendleriana* and *Pseudoroegneria spicata* seeds which were treated and planted in two differing terrains/substrates. Seeds were either non-treated (Control), incorporated into a pod (Pod), or incorporated into a primed-seed pod (P-Pod). Seeds were planted in either a ridgetop or hillslope soil collected from the West Side region of the Kaibab Plateau, USA.

| Species | Soils | Treatment | T<sub>50e</sub><sup>†</sup> | MET<sup>‡</sup> | FE<sup>β</sup> |
| --- | --- | --- | --- | --- | --- |
|  |  |  | -----days----- |  | % |
| *Poa fendleriana* | Ridgetop | Control | 17.6 (4.7) a* | 19.8 (4.3) a | 4.6 (1.9) b |
|  |  | Pod | 20.3 (2.4) a | 21.8 (1.7) a | 10.0 (3.0) b |
|  |  | P-Pod | 3.1 (0.7) b | 4.1 (1.0) b | 17.3 (5.2) a |
|  | Hillslope | Control | 13.6 (2.2) a | 16.9 (1.7) a | 10 (2.6) a |
|  |  | Pod | 15 (1.3) a | 17.8 (1.2) a | 8.6 (3.4) a |
|  |  | P-Pod | 4.6 (1.0) b | 5.3 (0.9) b | 18.6 (2.4) a |
| *Pseudoroegneria spicata* | Ridgetop | Control | 10.0 (2.9) a | 11.3 (2.7) a | 9.6 (3.7) b |
|  |  | Pod | 11.5 (2.1) a | 13.8 (1.4) a | 33.3 (5.8) a |
|  |  | P-Pod | 2.4 (0.5) b | 5.8 (0.8) b | 36.6 (4.3) a |
|  | Hillslope | Control | 8.2 (2.3) a | 9.7 (2.0) a | 13.3 (2.4) b |
|  |  | Pod | 7.9 (1.4) a | 9.2 (1.4) a | 18 (5.2) b |
|  |  | P-Pod | 2.7 (0.5) b | 4.7 (0.8) b | 38.6 (6.5) a |

<sup>†</sup>T<sub>50e</sub> = time to 50 percent emergence, <sup>‡</sup>MET = mean emergence time
<sup>β</sup>FE = final emergence
\*Significant differences ( $P \leq 0.05$ ) within a species and soil type
Numbers in parentheses represent s.e.m.

**Fig 2.**
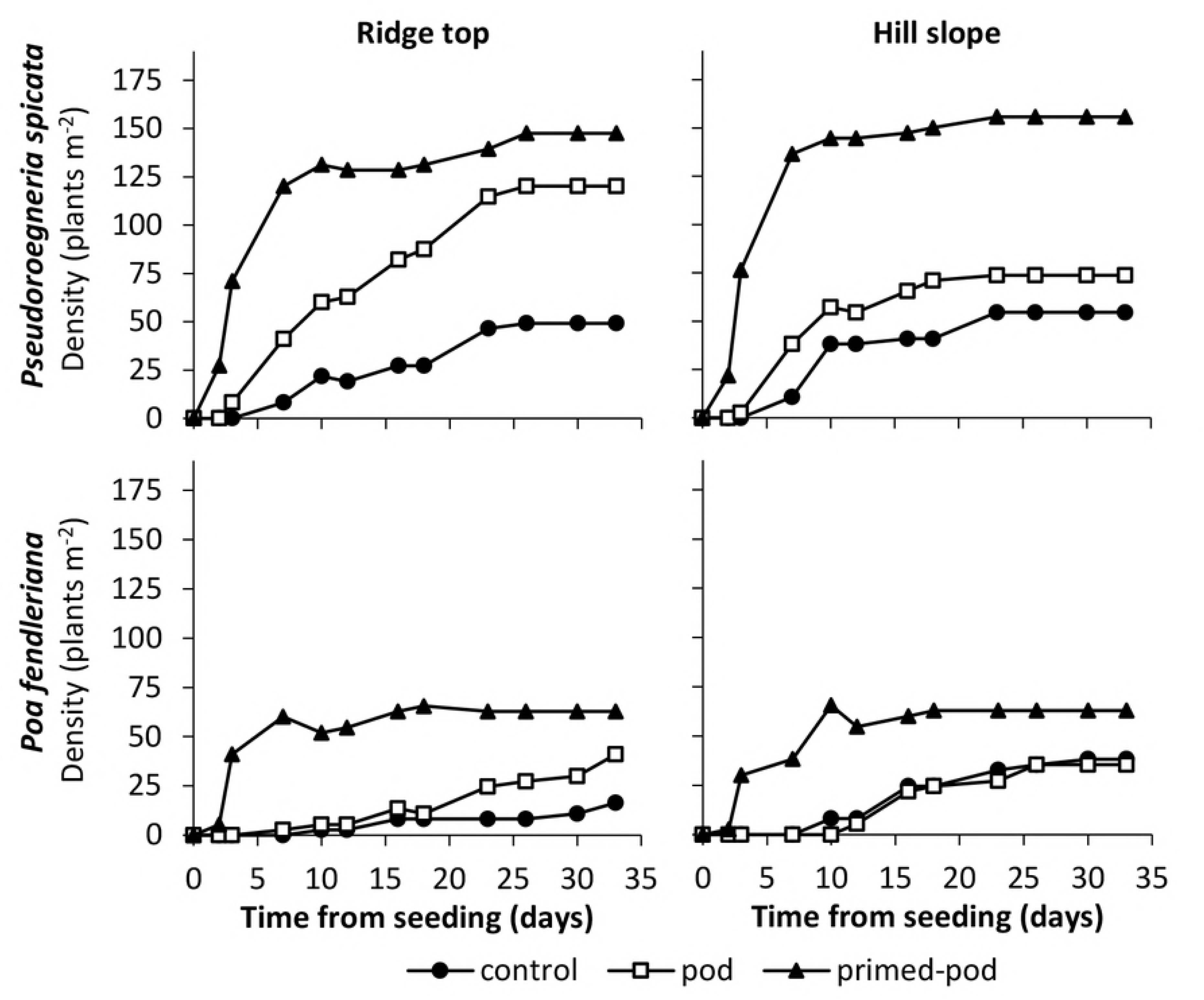
Plant Density (plants m^-2^) of two rangeland species (*P. spicata* and *P. fendleriana*) of ridgetop and hillslope soils. Readings taken from time of seeding until end of the study (day 35).

Final emergence (FE) was influenced by seed treatments for *P. fendleriana* in the ridgetop soil (*F* = 5.33, *P* = 0.03). By the conclusion of the study, *P. fendleriana* primed-seed pods in the ridgetop soil had produced 3.8 and 1.7-fold more seedlings than the control and non-primed-seed pods, respectively (Table 4). In the hillslope soil, despite *P. fendleriana* primed-seed pods showing early differences in seedling density over the other treatments, plant density was similar among all the treatments by the end of the study (*F* = 3.73, *P* = 0.11; Fig. 2; Table 4).

Seed treatments influenced final emergence for both soil types planted with *P. spicata* (ridgetop *F* = 22.21, *P* < 0.001, hillslope *F* = 6.41, *P* = 0.02). FE of *P. spicata* primed-seed pods were 2.9 and 3.8-fold higher than the control for ridgetop and hillslope soils, respectively. Non-primed-seed pods improved FE for *P. spicata* by 3.4-fold over the control in the ridgetop soil but was similar to the control in the hillslope soil (Table 4).

## Discussion

This study demonstrates that solid matrix priming (SMP) techniques can be an effective pre treatment to decrease seed germination time. Several studies have explored the use of priming to decrease seed germination timing of native dryland grass species [18, 26-28]. Unlike other proposed SMP techniques, this is the first study to demonstrate a procedure that does not require the seeds to be separated from the matrix after priming, which may reduce costs and improve efficiency of production. Rather, the seed and matrix material can be formed into a dough and extruded into pods or other shapes. The technology described in this study also requires less expensive production equipment, making it possible for small, localized processing. Small, local production would allow flexibility in application timing, which may reduce risks of possible seed deterioration due to extensive storage time [29]. Applying treated seeds as soon as possible is recommended for managers, though studies testing longevity of primed seed viability are needed.

In addition to seeds being enhanced through priming, planting seeds with SMP material in the form of extruded seed pods may also augment seeding success by improving the microsite surrounding the seed. The pod shape was designed to improve seed coverage by having a flat bottom and convex top. A broadcasted pod in this shape tumbling along the soil surface is more likely to come to rest with the bottom of the pod towards the ground [22]. It was also anticipated from our design that moisture would allow the SMP material in the pod to break down over the seeds, thus providing seed coverage and enhanced conditions for seed germination and growth. However, this study provides only moderate evidence that the SMP material used was beneficial at enhancing the microsite surrounding the seed. In only one of the four trials conducted, non-primed-seed pods showed improved emergence over non-treated seeds. In contrast, in all of the trials conducted, primed-seed pods exhibited faster emergence and a higher number of seedlings in comparison to non-treated seed. These results indicate that priming is the principal treatment responsible for primed-seed pods outperforming non-treated seeds.

One potential reason the non-primed seed pods did not show improved emergence under all the trials conducted, could be associated with an observation that seed pods disconnected with the soil surface as they dried. The super absorbent materials in the pods swelled with the addition of water, but when the pods dried, they would lift from the soil surface at the edges, which caused further and increased dry down within the pods. It is probable that seed pods and primed-seed pods could be improved by removing super-absorbent materials from the formulation to reduce the amount of swell-shrink in the pod and maintain the pod’s connectivity with the soil surface. Alternatively, the same ingredients used to form seed pods could be used to form pellets, which can be planted below the soil surface with a seed drill due to their shape. Under this scenario, swelling caused by the super-absorbent polymers may aid in seedling emergence by breaking through soil physical crust. Madsen et al. [13] demonstrated that extruded seed pellets made with super-absorbent polymers improved seedling emergence of *Artemisia tridentata* Nutt. ssp. *wyomingensis* Beetle & Young (Wyoming big sagebrush).

Primed extruded seed pods may improve restoration outcomes of autumn or spring plantings of native grasses by decreasing the time it takes for seeds to germinate and emerge from the soil. In the western United States, seeding practices typically occur in late autumn to early winter [30]. Between autumn and spring, significant seed loss and seedling mortality can occur. Long soil incubation times increase the probability for a predator [31] or pathogen attack on seeds [32]. Research by James et al. [6] and Boyd & James [33] indicates that over 70% of grass seeds planted in autumn can germinate prior to winter onset, but fail to emerge from the soil. Significant mortality may occur to these young, pre-emergent seedlings over the winter period as they are subjected to multiple prolonged freezing events, drought, pathogens, and expenditure of seed food resources [34]. Many of the invasive weeds, such as *B. tectorum* follow a different germination strategy by rapidly germinating and emerging from the soil after autumn rainfall. This species then grows to a size that can withstand harsh conditions of winter and is capable of quickly capturing available soil moisture and nutrient resources when growing conditions are suitable in winter and spring [14]. Primed-seed pods planted in autumn may mimic invasive annual grasses and experience reduced mortality due to rapid germination that allows the seedling to produce enough biomass to survive the harsh freezing conditions of winter.

Alternatively to avoid winter mortality, seeds could be planted in the spring; however, under this scenario, seeds may not germinate or germinate late in the season and subsequently not be sufficiently developed to survive through the summer drought period. Spring plantings of primed extruded seed pods may allow germination to occur early in the season. This would improve the probability that seminal roots of seedlings stay ahead of an advancing drying front and allow sufficient time for adventitious roots to develop before an extended drought period.

## Conclusions

The findings presented in this research provide a novel seed enhancement technology that decreases seed germination timing and improves seedling emergence for two cold desert grass species. Primed-seed pods may be beneficial in a variety of wildland and agricultural systems to improve direct seeding efforts. This technology is still in early stages of development and could be improved by future research that identifies materials that can not only be used for SMP, but also enhance the micro-environment within which seeds are planted.

## Acknowledgments

Funding for this research was provided by Kane and Two Mile Research and Stewardship Partnership. Oregon Department of Fish and Wildlife, US Fish and Wildlife Service Wildlife and Sport Fish Restoration Program, USDA Agriculture Research Service, and Brigham Young University.

## References

1. Nelleman C, Corcoran E. (Eds.). Dead Planet, Living Planet—Biodiversity and Ecosystem Restoration for Sustainable Development: A Rapid Response Assessment. United Nations Environment Programme. 2010.

2. D’Antonio CM, Vitousek PM. Biological invasions by exotic grasses, the grass/fire cycle, and global change. Annual Review of Ecology and Systematics. 1992; 23, 63–87. doi:10.1146/annurev.es.23.110192.000431

3. Svejcar T, Boyd C, Davies K, Hamerlynck E, Svejcar LN. Challenges and limitations to native species restoration in the Great Basin, USA. Plant Ecology. 2017; 218, 81–94. doi:10.1007/s11258-016-0648-z

4. Davies KW, Boyd CS, Beck JL, Bates JD, Svejcar TJ, Gregg MA. Saving the sagebrush sea: an ecosystem conservation plan for big sagebrush plant communities. Biological Conservation. 2011; 144, 2573–2584. doi:10.1016/j.biocon.2011.07.016

5. Hardegree SP, Jones TA, Roundy BA, Shaw NL, Monaco TA. Assessment of range planting as a conservation practice. Rangeland Ecology & Management. 2016; 69, 237–24 doi:10.1016/j.rama.2016.04.007

6. James JJ, Svejcar TJ, Rinella MJ. Demographic processes limiting seedling recruitment in aridland restoration. Journal of Applied Ecology. 2011; 48, 961–969. doi:10.1111/j.1365-2664.2011.02009.x

7. Bernstein EJ, Albano CM, Sisk TD, Crews TE, Rosenstock S. Establishing Cool-Season Grasses on a Degraded Arid Rangeland of the Colorado Plateau. Restoration Ecology. 2014; 22, 57–64. doi:10.1111/rec.12023

8. Taylor AG, Allen PS, Bennett MA, Bradford KJ. Seed enhancements. Seed Science Research. 1998; 8, 245–256. doi:10.1017/S0960258500004141

9. Madsen MD, Zvirzdin DL, Kostka SJ. Improving reseeding success after catastrophic wildfire with surfactant seed coating technology. ATSM International. 2013; 1569, 44–55. doi:10.1520/STP156920120181

10. Erickson TE, Shackelford N, Dixon KW, Turner SR, Merritt DJ. Overcoming physiological dormancy in seeds of Triodia (Poaceae) to improve restoration in the arid zone. Restoration Ecology. 2016; 24, S64–S76. doi:10.1111/rec.12357

11. Guzzomi AL, Erickson TE, Ling KY, Dixon KW, Merrit DJ. Flash flaming effectively removes appendages and improves the seed coating potential of grass florets. Restoration Ecology. 2016; 24, S98–S105. doi:10.1111/rec.12386

12. Madsen MD, Kostka SJ, Fidanza MA, Barney NS, Badrakh T, McMillan MF. Low-dose application of non-ionic alkyl terminated block copolymer surfactant enhances turfgrass seed germination and plant growth. HortTechnology. 2016; 26, 379–385.

13. Madsen MD, Hulet A, Phillips K, Staley JL, Davies KW, Svejcar TJ. Extruded seed pellets: A novel approach to enhancing sagebrush seedling emergence. Native Plant Journal. 2016; 17, 230–243. doi:10.3368/npj.17.3.230

14. Mack RN, Pyke DA. The demography of Bromus tectorum: variation in time and space. Journal of Ecology. 1983; 71, 69–93. doi:10.2307/2259964

15. Vaughn KJ, Young TP. Short-term priority over exotic annuals increases the initial density and longer-term cover of native perennial grasses. Ecological Applications. 2015; 25, 791–799. doi:10.1890/14-0922.1

16. Bradford KJ. Manipulation of seed water relations via osmotic priming to improve germination under stress conditions. HortScience. 1986; 21, 1105–1112.

17. Bewley JD. Seed germination and dormancy. Plant Cell. 1997; 9, 1055–1066. doi:10.1105/tpc.9.7.1055

18. Hardegree SP, Jones TA, Van Vactor SS. Variability in thermal response of primed and non-primed seeds of squirreltail [Elymus elymoides (Raf.) Swezey and Elymus multisetus (J.G. Smith) M. E. Jone]. Annals of Botany. 2002; 89, 311–319. doi:10.1093/aob/mcf043

19. Halmer P. Seed technology and seed enhancement. Acta Horticulturae. 2008; 771, 17–26. doi:10.17660/ActaHortic.2008.771.1

20. Taylor AG, Klein DE, Whitlow TH. SMP: solid matrix priming of seeds. Scientia Horticulturae. 1988; 37, 1–1. doi:10.1016/0304-4238(88)90146-X

21. Harman GE, Taylor AG. Improved seedling performance by integration of biological control methods at favorable pH levels with solid matrix priming. Phytopathology. 1988; 78, 520–525. doi:10.1094/Phyto-78-520

22. Madsen MD, Svejcar TJ. Development and application of “Seed Pillow” technology for overcoming the limiting factors controlling rangeland reseeding success. 2016; U.S Patent Application No. 14/039, 873, Patent Publication No. US9326451 B1.

23. Hardegree SP, Moffet CA, Roundy BA, Jones TA, Novak SJ, Clark PE, Pierson FB, Flerchinger GN. A comparison of cumulative-germination response of cheatgrass (Bromus tectorum L.) and five perennial bunchgrass species to simulated field-temperature regimes. Journal of Experimental Botany. 2010; 69, 320–327. doi:10.1016/j.envexpbot.2010.04.012

24. USDA. Terrestrial ecosystem survey of the Kaibab National Forest: Coconino County and Part of Yavapai County, Arizona. United States Department of Agriculture, Southwestern Region; 1984–89.

25. Cassel D, Nielsen D. Field capacity and available water capacity. In: Klute A, editor. Methods of soil analysis, part 2nd ed. Madison (WI): Soil Science Society of America; 1986. 1188 p.

26. Hardegree SP, Emmerich WE. Effect of Matric-Priming Duration and Priming Water Potential on Germination of Four Grasses. Journal of Experimental Botany. 1992; 43, 233–238. doi:10.1093/jxb/43.2.233

27. Hardegree SP. Matric priming increases germination rate of Great Basin native perennial grasses. Agronomy Journal. 1994; 86, 289–23. doi:10.2134/agronj1994.00021962008600020015x

28. Hardegree SP. Optimization of seed priming treatments to increase low-temperature germination rate. Journal of Range Management. 1996; 49, 87–92.

29. Tarquis AM, Bradford KJ. Prehydration and priming treatments that advance germination also increase the rate of deterioration of lettuce seeds. Journal of Experimental Botany. 1992; 43, 307–317.

30. Pyke DA, Chambers JC, Pellant M, Miller RF, Beck JL, Doescher PS, Roundy BA, Schupp EW, Knick ST, Brunson M, McIver JD. Restoration handbook for sagebrush steppe ecosystems with emphasis on greater sage-grouse habitat—Part Site level restoration decisions: U.S. Geological Survey Circular 1426, 62.2017. doi:10.3133/cir1426

31. St Clair SB, O’Connor R, Gill RA, McMillan BR. Biotic resistance and disturbance: rodent consumers regulate post-fire plant invasions and increase community diversity. Ecology. 2016; 97, 1700–1711. doi:10.1002/ecy.1391

32. Gornish ES, Aanderud ZT, Sheley RL, Rinella MJ, Svejcar TJ, Englund SD, James JJ. Altered snowfall and soil disturbance influence the early life stage transitions and recruitment of a native and invasive grass in a cold desert. Oecologia. 2015; 177, 595–606. doi:10.1007/s00442-014-3180-7

33. Boyd CS, James JJ. Variation in timing of planting influences bluebunch wheatgrass demography in an arid system. Rangeland Ecology & Management. 2013; 66, 117–126. doi:10.2111/REM-D-11-00217.1

34. Madsen MD, Davies KW, Boyd CS, Kerby JD, and Svejcar TJ. Emerging seed enhancement technologies for overcoming barriers to restoration. Restoration Ecology. 2016; 24, S77–S84.

